# Parasite-driven replacement of a sexual by a closely related asexual taxon in nature

**DOI:** 10.1101/143446

**Authors:** Jennifer N. Lohr, Christoph R. Haag

**Affiliations:** University of Fribourg, Department of Biology, Ecology and Evolution, Fribourg, Switzerland; Tvärminne Zoological Station, Hanko, Finland; Institute of Healthy Ageing, Department of Genetics, Evolution and Environment, University College London, London WC1E 6BT, UK.; Centre d’Ecologie Fonctionnelle et Evolutive, Université de Montpellier, France

**Keywords:** *Daphnia*, ecological replacement, reproductive mode, invasion biology

## Abstract

Asexual species are thought to suffer more from coevolving parasites than related sexuals. Yet, this prediction may be modulated by the fact that closely related sexuals and asexuals often differ in respects other than reproductive mode. Here, we follow the frequency dynamics of sexual and asexual *Daphnia pulex* in a natural pond that was initially dominated by sexuals. However, coinciding with an epidemic of a microsporidian parasite infecting both sexuals and asexuals, the pond was rapidly taken over by the initially rare asexuals. We experimentally confirm that asexuals are less susceptible and also suffer less from the parasite once infected. These results show the ecological replacement of a sexual taxon by a closely related asexual taxon, as driven by parasites. We suggest that this replacement is, however, not directly connected with the reproductive mode, but rather due to the recent introduction and invasive nature of the asexuals studied.

## Introduction

Theory on host-parasite interactions predicts that asexual species should suffer more from parasites compared to their sexual relatives. This is due to their lower genetic diversity and reduced ability to generate new allelic combinations, which are predicted to lead to a higher average parasite load (Jaenike 1978; Lloyd 1980; Hamilton 1980; Hamilton et al. 1990). This is thought to prevent asexuals from replacing sexuals within populations, because asexuals are thought to become increasingly parasitized as they become more common. The prediction that asexuals should be overparasitized has been investigated empirically in many study systems, with some studies finding evidence in support of the prediction (Lively & Jokela 2002; Kumpulainenet *et al*. 2004; Vergara *et al*. 2014) and others not (Hanley *et al*. 1995; Ben-Ami & Heller 2005; Elzinga *et al*. 2012).

While such variable results are not necessarily inconsistent with theory (asexuals are only predicted to be overparasitized on average, and not in every single case), they may also be due to differences between sexuals and asexuals, other than the reproductive mode. In fact, asexual taxa may only rarely differ from their closely related sexual relatives in reproductive mode alone (e.g. Lehto & Haag 2010; Gilabert *et al*. 2014; Kotusz *et al*. 2014). This is because the evolution of a new reproductive mode is often followed by secondary adaptations and differentiation, involving traits unrelated to reproductive mode (Meirmans *et al*. 2012; Gilabert *et al*. 2014). Furthermore, asexuals may have, from the outset, different genetic backgrounds, as they often arise via hybridization or polyploidy (Simon *et al*. 2003). These differences may add strong variation to the predictions of who should be more parasitized, and, in some cases, perhaps even lead to the inverse prediction.

One such case where asexuals might suffer less from parasites than sexuals is in the context of biological invasions. Asexuals are thought to be particularly prone to become invasive (Vandel 1928; Baker 1955; Peck *et al*. 1998; Haag & Ebert 2004), and indeed there is empirical data showing that asexuals are overrepresented among invasive species (Ellstrand & Schierenbeck 2000; Sakai *et al*. 2001). Additionally, invasive species do not share evolutionary history with the native parasite species in their invasive ranges. This can provide a parasite-mediated competitive advantage for invaders over residents, as local adaptation of parasites to resident hosts is thought to be common (Torchin *et al*. 2003). Hence, considering that asexuals are more common among invasive species and that parasites are likely more adapted to residents than to invaders, sexuals rather than asexuals may be predicted to suffer more from parasites during the early phases of invasion.

Here, we document the rapid replacement of a sexual by a closely related asexual taxon in nature and show experimentally that this replacement was very likely driven by parasites. We discuss the results with respect to sex-asex predictions, as well as with respect to invasion biology (the asexual taxon is invasive). The aim of this study is not to pinpoint the ultimate driver behind parasite-driven competition in sex-asex systems, but rather to showcase a specific empirical example where asexuals, rather than sexuals, enjoy a parasite-mediated advantage. We then discuss the implications of these findings for other empirical sex-asex comparisons.

Specifically, we studied cyclical and obligate parthenogenetic lineages of the freshwater cladoceran *Daphnia pulex* that occur in small rock pools to form dynamic metapopulations across the skerry archipelago of Southern Finland (Pajunen 1986; Pajunen & Pajunen 2003; Lehto & Haag 2010). In winter the ponds freeze to the bottom and only diapause stages survive. In our study area, about 5 to 10 % of the rock pools harbour *D. pulex*. Cyclical and obligate parthenogenetic *D. pulex* often inhabit different ponds, in part due to chance events caused by the dynamics of extinction and recolonization and in part due to different niche preferences with respect to water chemistry. In ponds with intermediate water chemistry, cyclical and obligate parthenogenetic populations coexist (Lehto & Haag 2010).

Obligate asexual lineages belong to the North American clade of *D. pulex* and have a history of introgression and contagious asexuality with cyclical parthenogens of the same clade, as well as the closely related *Daphnia pulicaria* (Innes & Hebert 1988; Tucker *et al*. 2013; Xu *et al*. 2015). It is unknown exactly when the asexuals were introduced to Europe, but it was probably near the beginning of the 20^th^ century in the ballast water of transport ships returning from the Laurentian Great Lakes, a transport route common to many aquatic invasive species (Carlton 1985; Williams *et al*. 1988). The cyclical parthenogenetic *D. pulex* in our study system belong to the European clade, which, however, may represent a different species (Mergeay *et al*. 2008; Markova *et al*. 2013). In the absence of a formal description of the two taxa as different species, we here still use the species name *D. pulex* for both, but refer to them as closely related taxa. Obligate asexuals (hereafter “asexuals”) produce both live-born offspring and diapausing stages asexually, whereas cyclical parthenogens (hereafter “sexuals”) have several asexual generations (production of live-born offspring) and typically one sexual generation (production of diapause stages) per year (Innes & Hebert 1988; Paland *et al*. 2005; Lynch *et al*. 2008; Heier & Dudycha 2009). Asexuals are genetically highly uniform, whereas sexuals show moderate levels of genetic variation, typical of other cyclical parthenogenetic *Daphnia* species (Ward *et al*. 1994; Haag *et al*. 2005; Walser & Haag 2012).

Here we followed the frequencies of sexuals and asexuals in one such mixed natural pond over 14 years (1999-2013). In 2006, a strong epidemic caused by a highly virulent parasite, *Gurleya vavrai*, coincided with the total replacement of formerly dominating sexuals by obligate asexuals. G. *vavrai* is an endemic microsporidian parasite in Europe (Green 1974; Refardt *et al*. 2002), with no record of the parasite occurring in North America. Infections are localized in the epidermis and cause a progressive whitening of the body as the infection spreads, with high virulence (Friedrich *et al*. 1996; Stirnadel & Ebert 1997; Little & Ebert 1999). Using a series of field and laboratory experiments, we assessed the relative competitiveness of sexuals and asexuals in the presence and absence of the parasite, the susceptibility of sexual and asexuals hosts, as well as life history traits of infected and uninfected individuals. The goal of these experiments was to investigate whether the reversal in relative abundance observed in the natural pond was likely caused by parasites.

## Materials and Methods

### Monitoring of the natural pond

One pond, SK-39 (59°49’55.7”N 23°15’17.6”E, pond surface 1.6 m^2^, depth 0.3 m) on the island of Skallothomen, which contained both sexuals and asexuals in 1999, was followed for the present study because it was initially uninfected by *G. vavrai*, but became infected in 2006. In 2000 and again in 2008, no animals were observed in the pond during routine visits, possibly because appropriate conditions for the hatching of diapause stages did not occur in those years. In all other years, namely 1999 to 2015, *D. pulex* were present. We monitored the frequencies of sexuals and asexuals in a total of 26 samples taken between 1999 and 2013 (total *N* = 1848 individuals, for exact sampling dates and sample sizes per date, see data deposited on the Dryad digital repository). The breeding type (sexual, asexual) of each individual in these samples was assessed using cellulose-acetate electrophoresis (Hebert & Beaton 1993) at the PGI locus (phosphoglucose isomerase, EC 5.3.1.9.), for which the sexuals and asexuals have consistently different genotypes (Lehto & Haag 2010). On July 14^th^ 2006, we noted a strong *G. vavrai* infection in our samples. Even though previous samples were not systematically checked for this parasite, a previous epidemic would not have gone unnoticed. We subsequently took additional samples to estimate the parasite prevalence in sexuals and asexuals (20, 25, and 30 July, 29 September). Prevalence was assessed by visually inspecting the carapace color with light from the top and against a dark background. Checking for a white carapace detects only late stage infections, while early stage infections are pooled with the uninfected animals. Thus, all prevalence estimates are underestimates. From 2007-2013 we recorded only presence/absence of infections.

#### Outdoor experiment: Origin and handling of hosts and parasites

Ten ponds composed purely of sexuals and ten ponds of purely asexuals were sampled in the Tvärminne area in May 2010 (Table 1). From each of these ponds, we randomly sampled 100 individuals to capture a representation of the genetic variation present. These 20 geographically separate ponds should be representative of ponds across the study area: Hence the experiment was designed to test whether an average, uninfected asexual in this metapopulation suffers less from infection (and/or is less susceptible) than an average, uninfected sexual in this metapopulation. From each of these 20 populations, an outdoor culture was established in buckets, and left outdoors under ambient conditions on the island of Furuskär (59°49’58”N 23°15’49”E). Before introducing the *Daphnia*, each bucket was filled with 40L of 0.02 mm-double filtered water from a nearby pond, not colonized by *Daphnia*. The 100 individuals were then left for two months to reproduce and increase in number. The only exception was a bucket of asexuals from SK-39, which had already been started using a single, uninfected individual in 2007. The breeding type (sexual, asexual) of each bucket was confirmed using the PGI locus as described above. In July of 2010 these cultures were harvested and used as the experimental animals for the outdoor experiment. Individuals were checked for parasite infection to ensure the absence of any infected individuals at the start of the experiment. Four additional cultures of *D. pulex*, which had been maintained in buckets on the island of Furuskär from 2007 to 2010 with *G. vavrai* infections, were used as the source material for the spore cocktail prepared for infections. In two of these cultures, *G. vavrai* was grown on sexuals, and in the two others on asexuals (Table S1). The two cultures per host taxon were combined to produce the spore cocktails for infection (one cocktail derived from sexual hosts, one from asexual hosts).

#### Outdoor experiment: Experimental set-up

We released 100 sexuals and 100 asexuals, obtained as described above, into each of 20 new outdoor cultures (40L bucket cultures). This resulted in ten experimental pairings (each pair consisting of sexuals from one sexual pond and asexuals from one asexual pond, Table S1), with each pair repeated twice. One replicate per pair was exposed to *G. vavrai* spores and the second served as the control culture without parasite exposure. *D. pulex* infected with *G. vavrai* were ground up (individuals from all four cultures pooled) and distributed equally across the ten infection replicates. In the control treatment, a comparable amount of ground-up, uninfected *D. pulex* were added as a placebo.

#### Outdoor experiment: Recorded parameters

The cultures were left in the field, and samples were obtained from each culture at three time points: August 2010, September 2010 and October 2010. At each sampling event, 50 individuals were removed per culture and checked visually for infection. Subsequently, the same 50 individuals were typed as sexual or asexual using the PGI locus. In this way the prevalence of infection in sexuals versus asexuals could be monitored over the course of the experiment. Note that we were mainly interested in whether the frequency changes differed between treatments. The overall frequency changes are difficult to interpret because they are influenced by water chemistry (Lehto & Haag 2010). Note also that these estimates do not include potential differences in the production of diapause stages.

#### Laboratory experiment: Origin and handling of hosts and parasites

In June 2013, *D. pulex* were collected from various ponds in the Tvärminne area. Both sexuals and asexuals were collected, as well as *G. vavrai*-infected individuals of both taxa (Table S1). All individuals were transported live to the laboratory in Switzerland for immediate use. In the laboratory, one clonal line (lines started by single females and maintained exclusively by clonal reproduction in both the sexuals and asexuals) was established from each of the pond samples. Each line was propagated in a 400 ml glass jar filled with a water medium designed for *Daphnia* culturing (ADaM; Klüttgen *et al*. 1994) and fed with unicellular green algae (*Scenedesmus obliquitus*) *ad libitum* under a summer photo-period of 16:8 light:dark and a temperature of 20 °C. The lines were maintained in this way for two weeks, and their breeding type was confirmed using the PGI locus.

Two separate *G. vavrai* cultures were established in the laboratory: one obtained from and grown on sexuals and a second obtained from and grown on asexuals. These parasite types are referred to as sexual and asexual *G. vavrai* isolates, respectively, referring to their host type. In both cases the *D. pulex* hosts on which the spores were grown were those on which the parasite was collected, which were different clones from those used in the experiments (Table S1).

#### Laboratory experiment: Experimental procedures

For the infection experiment, we used eight clonal lines (four sexual lines and four asexual lines) with 30 replicate individuals per line and each line originating from a different pond (Fig. S1; Table S1). We started by isolating 50 individuals from each of the clonal lines (each placed individually in a 50 ml falcon tube) to ensure we would have at least 30 individuals at the beginning of the experiment. The animals were fed daily with 2,500 cells of the algae *S. obliquitus* and were kept in a climate chamber with a photo-period of 16:8 light:dark and a temperature of 20 °C. The ADaM culture medium was changed three times per week (Monday, Wednesday and Friday). The animals were passed through three generations under these pre-experimental conditions in order to remove maternal effects. Day zero of the experiment was when the fourth-generation offspring (third-clutch offspring of the third-generation females) were isolated into new tubes.

For infection, two separate spore cocktails were prepared: one from ground-up sexual hosts infected with *G. vavrai* and another from ground up infected asexual hosts infected with *G. vavrai*. Infected hosts came from two separate ponds for both the sexual and asexual cultures (Table S1). The total number of spores was equalized between the two treatments (spore numbers were estimated using a Neubauer improved counting chamber) and then distributed equally over the respective tubes. This resulted in approximately 60,000 *G. vavrai* spores being added to each tube in the parasite treatments. For the control treatment, a cocktail of ground up uninfected *D. pulex* was added to the tubes.

Offspring from 30 mothers per clone (and eight clones: four sexual, four asexual) were used as the experimental animals, resulting in a cohort of 240 animals at the start of the experiment. Each clonal line then received three different treatments, with ten replicates per treatment: (1) 60,000 spores of *G. vavrai* from asexual hosts, (2) 60,000 spores of *G. vavrai* from sexual hosts, (3) ground-up, uninfected *D. pulex* (control).

#### Laboratory experiment: Recorded parameters

Recorded parameters from the experiment were age at first reproduction, reproductive output (total number of offspring recorded across all changing events), age at death, as well as infection status and number of spores at death. To determine the number of spores at death, each individual was homogenized in 0.3 ml of medium and the concentration of spores was determined using a Neubauer Improved counting chamber.

### Data analysis

All statistical analyses were performed using the software R (R Core team 2013). For the outdoor competition experiment, we used a generalized linear-mixed model with a binomial error distribution to look at differences in the frequency of the sexuals vs. asexuals (R program lme4). For data analyses, each sexual individual was coded as 1, and each asexual individual as 0. We used treatment (control, parasite-exposed) and sampling time point as fixed factors, whereas replicate culture and pond pair were treated as random factors, with replicate culture nested within treatment: breedingtype ~ treatment*time + (1|treatment/replicate) + (1|pair of ponds), family = binomial. We used a similar generalized linear mixed model to investigate differences in parasite prevalence (proportion of individuals infected, each individual being counted as either infected or uninfected) between the sexual and asexual cultures, but here we used the parasite-exposed treatment only: infection_status ~ breedingtype*time + (1|replicate), family = binomial. Before running the analysis, we verified model assumptions and absence of over-dispersion by calculating the sum of the squares of the Pearson residuals and comparing them with the residual degrees of freedom using a chi-squared test.

Data on age at death, reproductive output and age at first reproduction from the laboratory experiment were evaluated using linear mixed models in R, with the nlme package. Breeding type (asexual or asexual), and treatment (control, infected) were set as fixed factors and clone as a random factor: trait ~ breedingtype*treatment, random= ~ (1|breedingtype/clone). Only those individuals that successfully became infected were used for comparison between treatments. We then used the same analysis exclusively on the infection treatment (and those individuals that actually became infected), but now used spore origin instead of treatment to test whether the different life history traits were affected by whether the *G. vavrai* spores were derived from sexual or asexual host clones. Using the same data, we also tested for differences in spore load at death using the same model as for the life history traits, as well as for differences in parasite prevalence (number of exposed individuals that became infected). For the latter, we used a generalized linear mixed model, with infection success as the response variable (1 = successfully infected, 0 = infection failed) and a binomial error distribution. Breeding type and spore type (sexual or asexual host origin) were held as fixed factors, whereas clone was a random factor nested within breeding system: infection status ~ breedingtype*sporetype, random= ~ 1|breedingtype/clone, family = binomial. As above, we verified model assumption and absence of over-dispersion before running the analyses.

## Results

### Dynamics in the natural pond

There were strong seasonal dynamics in the relative frequencies of sexuals and asexuals (Fig. 1). Asexuals were more frequent early in the season, and sexuals dominated in late season. Yet, across years, the starting frequencies of sexuals increased (reaching 93 % on June 1^st^ 2006), as the pond became more and more dominated by sexuals. By the end of June 2006, the frequency of sexuals had increased to 98 %. Then, on July 14^th^ 2006 we detected many individuals (sexuals and asexuals not distinguished) with *Gurleya vavrai* infections, and at the same time a dramatic decrease in the frequency of sexuals to 41 %. This decline continued over the course of the growing season down to 0 % (9 % on July 20^th^, 4 % on July 25^th^, 2 % on July 30^th^, and 0 % on September 28^th^). In the following years (two samples in 2007, and one in each of 2010, 2012, and 2013), we found only asexuals (Fig. 1).

**Figure 1.**
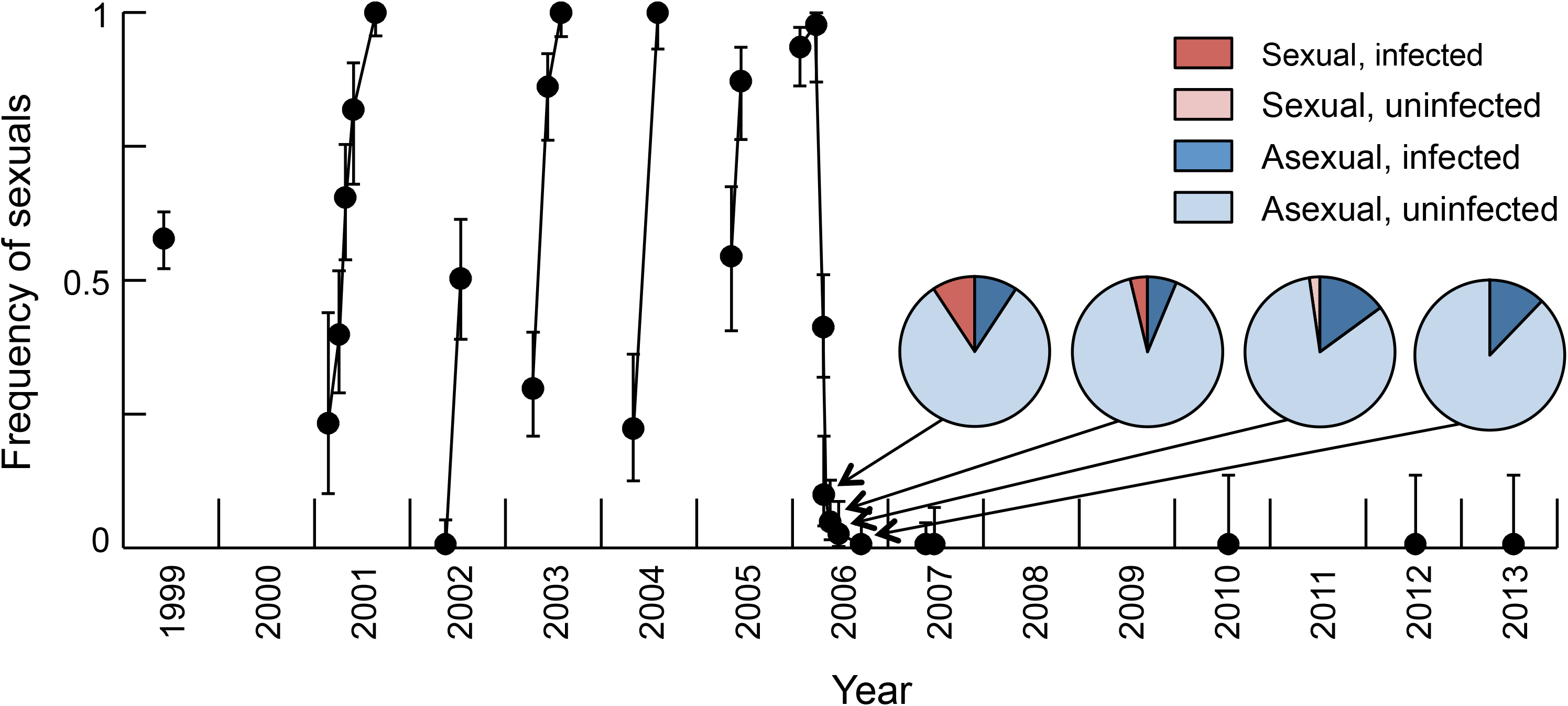
Frequency of sexuals in pond SK-39. Samples taken in the same year are connected with a line. The error bars correspond to 95 % confidence intervals according to the modified Wald method (Agresti & Coull 1998). The four pie charts represent data on prevalence whenever systematically recorded (red: sexuals, blue: asexuals, dark: infected, light: uninfected).

No quantitative measure of prevalence was recorded on July 14^th^ 2006, but we noted many infected individuals. On July 20^th^, many dead, infected individuals were found in the pond, the breeding type of which could not be assessed. Among those living, we found 10 infected individuals in a sample of 54 (prevalence = 19 %). Of the 10 infected individuals, 5 were sexuals and 5 were asexuals, whereas all of the uninfected individuals were asexuals (Fig. 1; Fisher’s exact test, *P* < 0.0001). Prevalence then rapidly dropped to low levels (2 % on July 25^th^, 4 % on July 30^th^, and 7 % on September 28^th^). The higher prevalence in sexuals was still significant on July 25^th^ (infected: 3 sexuals, 5 asexuals, uninfected: no sexuals, 72 asexuals, *P* = 0.0007), but not anymore on July 30^th^, when all 13 infected individuals were asexuals, whereas 2 among 74 uninfected individuals were sexuals (*P* > 0.5). The latter result shows that not all sexuals became infected and died during the epidemic. From 2007-2013, when only asexuals were left in the pond, we recorded infected individuals in all samples, though the infections were never as clearly abundant as during the beginning of the epidemic.

### Outdoor experiment

Asexuals performed relatively better in the competition experiment in the presence of the parasite *G. vavrai* than in their absence (Fig. 2). The infected cultures had higher frequencies of asexuals than the control cultures (*z* = -3.04, *df =2, P* = 0.002). This was true at both sampling time points and thus sampling time was not a significant factor in the linear mixed model (*z* = 0.60, *df =2, P* = 0.551). In addition, in the parasite treatment, a greater number of sexuals became infected, as opposed to asexuals (*z* = 2.21, *df =2, P* = 0.038; Fig. 2). Again sampling time point was not significant (*z* = 0.41, *df =2, P* = 0.389).

**Figure 2.**
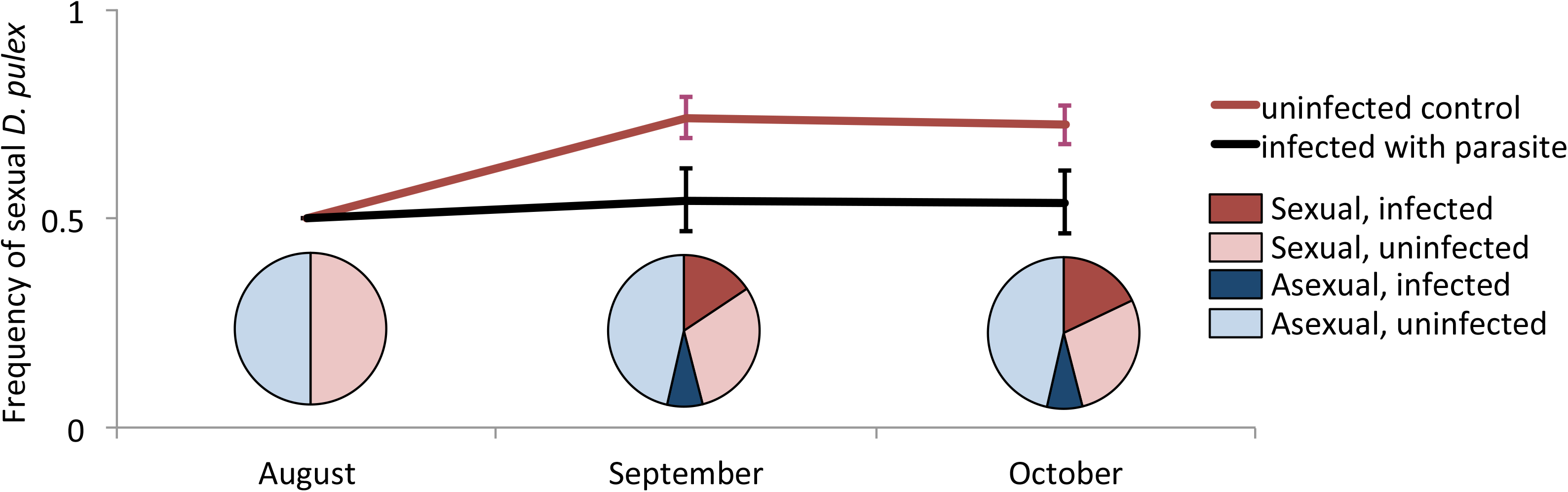
Outcome of the competition experiment between asexual and sexual *Daphnia pulex* exposed to *G. vavrai* in the summer of 2010. In August, the starting point of the experiment, there were an equal number of sexual and asexual individuals. Line graphs show the change in the frequency of sexuals and asexuals over the three month experiment. Error bars represent standard errors of the means across ten replicates. Pie charts show the proportion of the sexuals and asexuals infected with *G. vavrai* (red: sexuals, blue: asexuals, dark: infected, light: uninfected).

### Laboratory experiment

In line with the results from the outdoor experiment, 76.3 % of the sexual individuals exposed to *G. vavrai* spores became infected, whereas only 25.0 % of the asexuals did so (*z* = 4.96, *df* = 2, *P* < 0.001; Fig. 3a). The comparison of the infected individuals (spore origins not distinguished) with the uninfected controls revealed clear negative fitness effects of infection: infected individuals died sooner (*t* = -5.14, *df* = 149, *P* < 0.001) and were less fecund (*t* = -5.90, *df* = 149, *P* < 0.001), while there was no significant effect for age at maturity (*t* = 1.47, *df* = 149, *P* = 0.143; Fig. 4). In no case was breeding system a significant factor on its own, suggesting that there was no clear main difference in life-history traits between sexuals and asexuals under the experimental conditions (age at death: *t* = 0.52, *df*= 6, *P* = 0.620; age at maturity: *t* = 0.04, *df*= 6, *P* = 0.972; reproductive output: *t* = 1.60, *df* = 6, *P* = 0.163). However, there was a significant interaction between breeding type and infection for reproductive output (*t* = -3.10, *df* = 149, *P* = 0.002), suggesting that infection reduced the reproductive output of sexuals more strongly than that of asexuals, whereas asexuals had a somewhat lower reproductive output in the controls. The interaction was non-significant for age at death (*t* = -1.02, *df*= 149, *P* = 0.307) and for age at maturity (*t* = -0.51, *df*= 149, *P* = 0.614).

**Figure 3.**
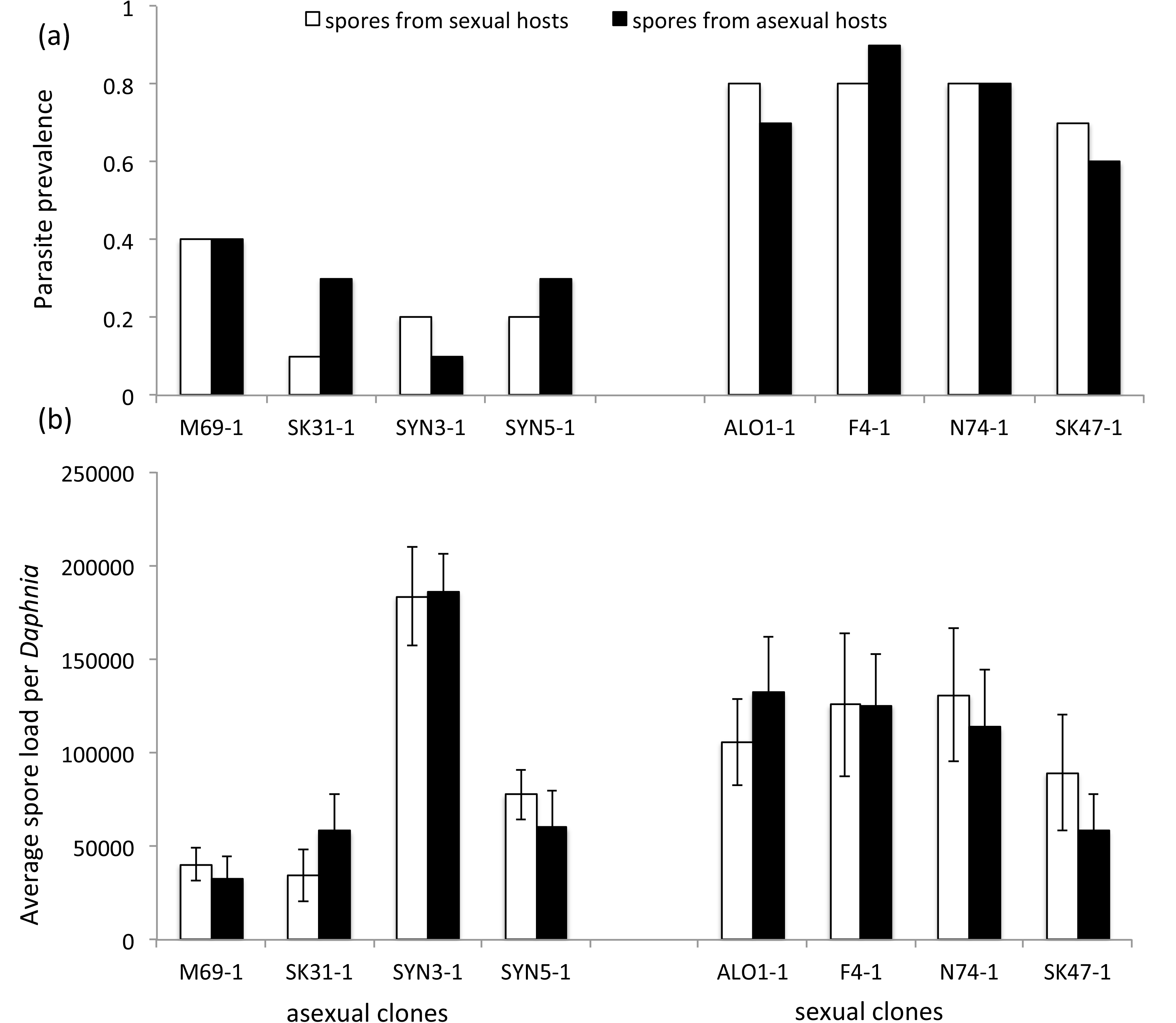
(a) Spore load at death and (b) parasite prevalence for the four asexual and four sexual clones used in the laboratory experiment. Error bars show the standard error of the mean values.

**Figure 4.**
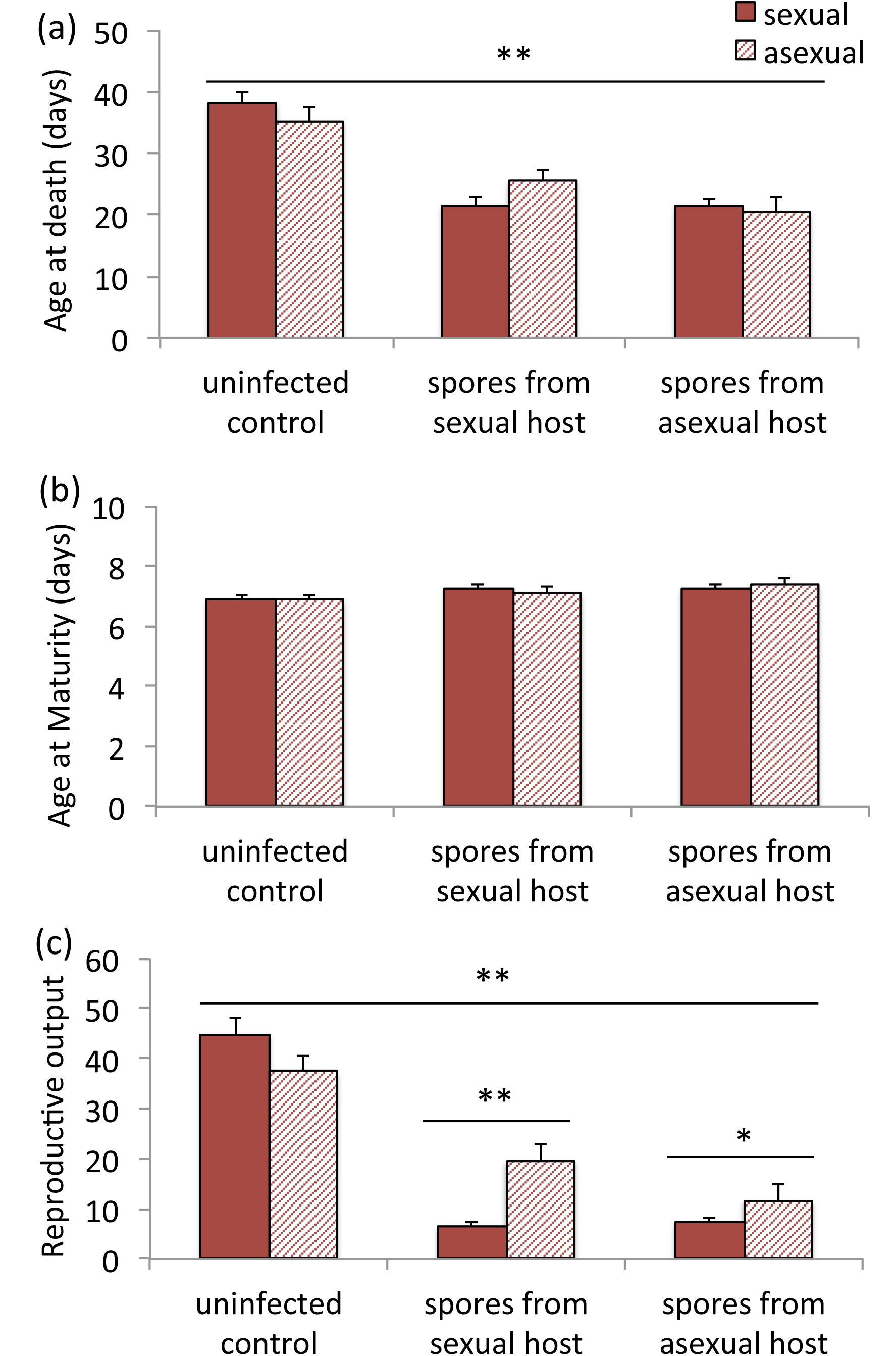
Life-history traits of asexual and sexual *Daphnia*, from either the control treatment, infection with spores of asexual origin treatment, or infection with spores of sexual origin treatment for (a) age at death, (b) age at first reproduction and (c) reproductive output (number of offspring per mother). Error bars show the standard error of the mean values. *P < 0.05, **P < 0.005.

Looking only at the infected individuals, there was no significant difference in spore load at death between sexual and asexual *Daphnia* (*t* = -0.60, *df* = 71, *P* = 0.553; Fig. 3b). Furthermore, the spore origin did not appear to affect infection success or spore load, either overall or differently between sexuals and asexuals (non-significant main effects: infection success: *z* = 1.03, *df* = 2, *P* = 0.304; spore load at death: *t* = 0.20, *df* = 150, *P* = 0.845; and non-significant interactions with breeding type: infection success: *z* = -1.28, *df=* 2, *P* = 0.200; spore load at death: *t* = 0.64, *df*= 71, *P* = 0.522). However, spore origin affected the reproductive output. Spores obtained from asexual hosts reduced the reproductive output of asexuals more than spores obtained from sexual hosts, whereas no such effect was observed in the sexuals (breeding type x spore type interaction: *t* = -2.85, *df* = 71, *P* = 0.006). Finally, spore origin did not affect age at death, or age at maturity (breedingtype*sporetype interaction; age at death: *t* = -1.51, *df*= 71, *P* = 0.135; age at maturity: *t* = -0.25, *df* = 71, *P* = 0.805; Fig. 4).

## Discussion

This study documents the replacement of an initially dominating sexual taxon by an initially rare asexual taxon in nature. This replacement did not occur gradually, as would be expected if the asexuals had an overall higher fitness. Rather, the replacement happened rapidly and was tightly linked with an epidemic caused by a virulent parasite, which infected both sexuals and asexuals. Subsequent field and laboratory experiments strongly support a causal role of the parasite in this replacement: Asexuals were less susceptible to infection, suffered less from infection than sexuals, and their relative performance in a competition experiment was enhanced in the presence of parasites. Independent of the reproductive mode of the two competitors, this is an example showing that parasites can strongly alter interactions between closely related taxa. Parasites have been implicated in ecological replacements between closely related species several times before, including in the replacement of residents by invaders (Thomas *et al*. 2005; Hatcher *et al*. 2006; Hatcher *et al*. 2008). Though, to our knowledge, in none of these previous cases could the replacement be monitored so closely in nature, or coupled with experimental evidence for a causal role of the parasite.

Our results also show that parasites can lead to rapid changes in the ecological frequency dynamics of coexisting sexual and asexual taxa. Discussion on competition between sexuals and asexuals with regards to parasites is often framed within Red Queen dynamics, whereby asexuals are predicted to become overinfected with parasites (at least on average), as they cannot evolve as fast as their sexual competitors (Jaenike 1978; Lloyd 1980; Hamilton 1980; Hamilton *et al*. 1990). In contrast, here we find that asexuals replace sexuals (and not the other way around) as a consequence of a parasite-driven advantage.

While our data are not suitable to pinpoint the ultimate cause behind this parasite-mediated advantage of asexuals, it seems likely that factors other than reproductive mode have played an important role. First, due to the invasive nature of the asexuals in this region, they do not share a long evolutionary history with the parasite (Innes & Hebert 1988; Ward *et al*. 1994). The parasite is endemic to Europe (Green 1974; Friedrich *et al*. 1996) and may thus not have had enough time to adapt to the asexuals. Second (and perhaps also due to their recent invasion of the region), asexuals are relatively rare in Southern Finland compared to sexuals (Lehto & Haag 2010). This is important because theory on host-parasite coevolution predicts that parasites should specialize on the most common genotypes (Jaenike 1978; Decaestecker *et al*. 2007: Salathé *et al*. 2008). However, the advantage of asexuals in the presence of parasites may only be transitory: once asexuals become abundant, parasites are predicted to adapt to them (Morran *et al*. 2011). Indeed, in our laboratory experiment, the spores obtained from asexual hosts were more virulent to asexuals than spores from sexual hosts. While this test was not replicated (we only tested one mixture of two parasite isolates from each of the two host types), this result suggests that some effects of parasite adaptation towards asexual hosts may have started to become visible. Finally, we cannot entirely exclude that the higher fitness of the asexuals in the presence of the parasite is due to the adaptation of asexuals to the parasite (rather than of the parasite to sexual hosts). However, this seems less plausible given the likely lack of a shared long-term evolutionary history.

As outlined in the introduction, it is also possible that our results are indirectly linked to reproductive mode. Asexuals are overrepresented among invasive species (Ellstrand & Schierenbeck 2000; Sakai *et al*. 2001), and obligate asexual *Daphnia* have been shown to be effective invaders who have rapidly replaced resident populations of closely related sexuals in other parts of the world, not only in Finland (Mergeay *et al*. 2006; Fadda *et al*. 2011; Duggan *et al*. 2012; So *et al*. 2015). Even though reproductive assurance (Baker 1955) does not differ between sexual and asexual *Daphnia* (due to cyclical parthenogenesis, also a single sexual individual can establish a population on its own), the success of obligate asexual *Daphnia* as invaders may still be linked to reproductive mode (though sexual *Daphnia* species can be invasive too; see Searle *et al*. 2016). First, obligate asexuality may allow a particularly successful invasive genotype to be “frozen” and thus shielded from segregation and mixing with other genotypes. In fact, what is essentially a single clone of *D. pulex* (or a group of closely related clones) is apparently responsible for the invasion of freshwater habitats in Africa and Southern Europe (Mergeay *et al*. 2006; Fadda *et al*. 2011). Second, colonization by a single cyclical parthenogenetic individual leads to within-clone mating during the production of diapause stages, and within-clone mating is known to lead to strong inbreeding depression in this and other *Daphnia* species (Deng & Lynch 1997; Lohr & Haag 2015), also specifically with regards to parasites (Haag et al. 2003).

Thus, our results suggest that parasites may have played an important role in the rapid establishment of these obligate asexual invaders, perhaps not only in Southern Finland but also elsewhere. In stark contrast, in the native range of asexual *D. pulex*, they are mostly outcompeted by sexuals if the two co-occur locally (Innes & Gin 2014). Though the possible involvement of parasites in the latter pattern is not known, this is potentially consistent with the “enemy release” advantage for invasive species, whereby invaders leave behind natural enemies from their native range and suffer less from newly encountered enemies in their introduced range (Keane & Crawley 2002; Torchin & Mitchell 2004).

Our study shows that asexuals do not always suffer more from parasites than sexuals. On the contrary, under certain circumstances, such as in the invasion context studied here, asexual can benefit from parasitism. Thus, the ecological context can modulate the generally predicted patterns of parasitism in sexual versus asexual species. This highlights an important limitation in the interpretation of empirical comparisons between closely related sexual and asexual species. These types of sex-asex comparisons are often used to investigate hypotheses related to the maintenance of sexual reproduction (Otto & Lenormand 2002), and while such comparisons can offer key insights, sexuals and asexuals often differ in many traits other than just the reproductive mode (e.g. Lehto & Haag 2010; Gilabert *et al*. 2014; Kotusz *et al*. 2014). This study highlights that it should not be assumed that patterns found between sexual and asexual species are driven solely by the reproductive mode.

## Acknowledgements

We thank R. Schleberger, A. Marcelino, C. Reisser, D. Fasel, D. Frey, M. Lehto, C. Stritt, and E. Hürlimann for help in the field and in the laboratory. We are very grateful to the staff of the Tvärminne Zoological Station of Helsinki University for help and support during fieldwork. We thank D. Ebert, J. Mergeay and three anonymous reviewers for their helpful comments. This work was supported by the Swiss National Science Foundation (Grant no. 31003A_138203), the University of Fribourg, and the European Union (Marie Curie Career Integration Grant PCIG13-GA-2013-618961, DamaNMP).

## Tables – supplementary material

**Table S1:**
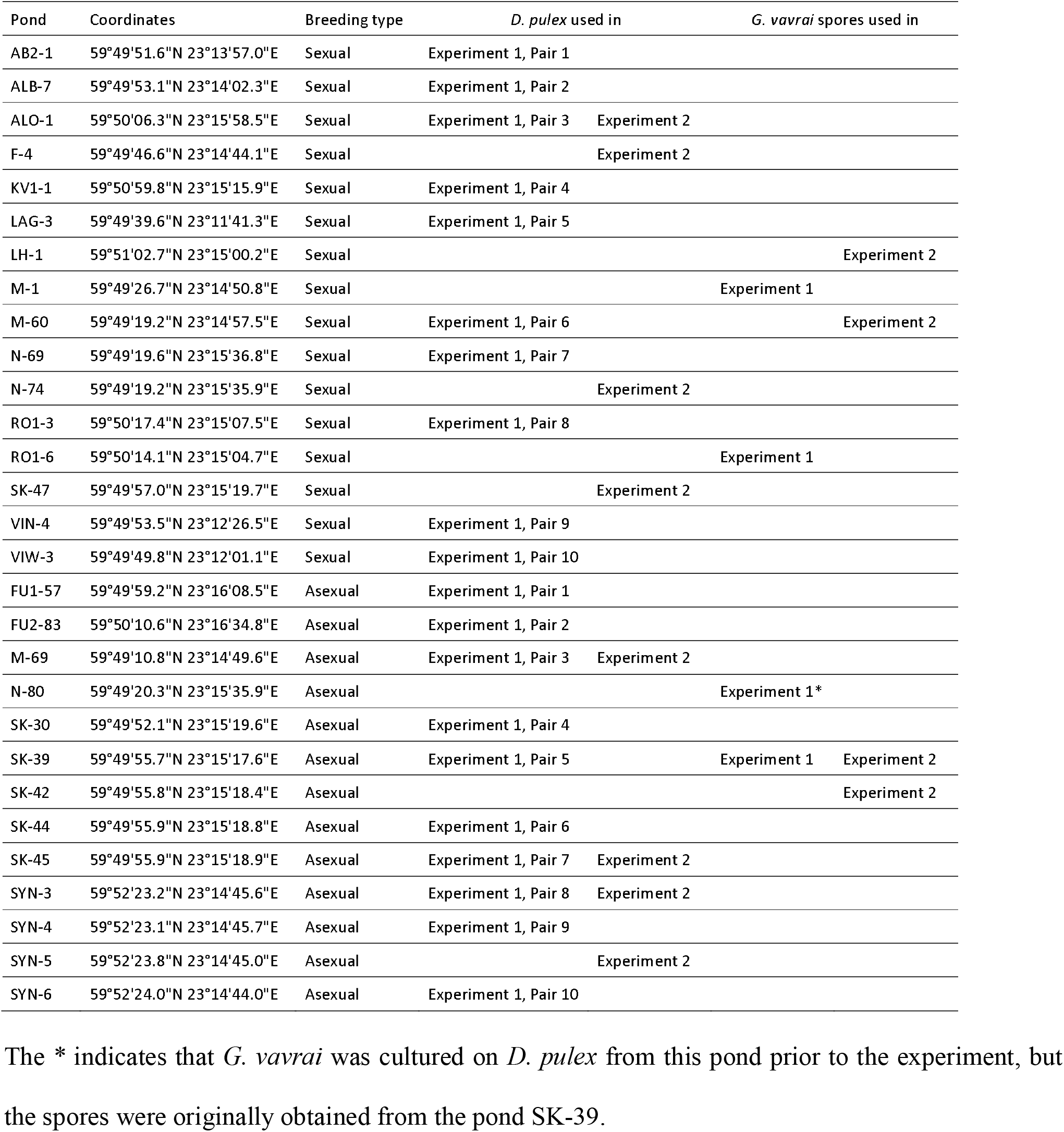
Host and parasite origins for the outdoor experiment (experiment 1) and the laboratory experiment (experiment 2). Ponds used both for *G. vavrai* spores in experiment 2 and for *D. pulex* in experiment 1 were previously uninfected (an uninfected *Daphnia* sample was obtained from these ponds for experiment 1).

Figure S1. Experimental design of the laboratory experiment. Four sexual and four asexual clones were each established from an independent pond within the Finnish archipelago.

